# Many are called but few are chosen – Multiple clonal origins greatly elevate the functional heterogeneity of tumors

**DOI:** 10.1101/2020.09.01.277848

**Authors:** Bingjie Chen, Xianrui Wu, Yongsen Ruan, Yulin Zhang, HJ Wen, Ping Lan, Chung-I Wu

## Abstract

Each tumor is usually accepted to be of a single origin from a progenitor cell. The shared evolutionary paths impose a limit on the nature of genetic diversity of the tumor. However, there are also numerous stem cell niches with independent proliferation potentials. To reconcile the contrasting perspectives, we propose a model whereby each tumor is of multiple clonal origins but the most proliferative one would eclipse other minor clones. The detection of the minor clones would entail an extreme scheme of large-number but small-volume sampling. In two cases of colon tumors so sampled, one indeed has 13 independent clones of disparate sizes and even the smaller clones have tens of thousands of cells dispersed non-locally. The other, much larger, tumor has only one prevailing clone that engulfs two tiny patches of minor clones. In both cases, the expanding clone spawns a hierarchy of subclones that resemble vassal states on its wake of expansion. The timing of metastasis can also be mapped to the precise stage of the clonal expansion. In conclusion, multiple independent clones, likely common but difficult to detect, can greatly elevate the non-neutral diversity within a tumor. This much-elevated diversity has many theoretical and clinical implications.

## Introduction

While it has been widely accepted that a tumor, or tumors, of an individual usually evolve from a single progenitor cell (Nowell 1976), this single-origin view raises many questions. As Cairns (Cairns 1975) pointed out in his seminal paper, each multi-cellular organism is compartmentalized into numerous small units, usually in the form of stem-cell niches. Each unit is an independently evolving entity and, once in a while, one of the units would evolve into an out-of-control proliferative population, manifested clinically as tumor.

As an evolutionary unit, each niche harbors stem cells with a population size of N. Functional studies often arrive at an estimate of N < 100 (Snippert, et al. 2010; Baker, et al. 2014). Recently, analysis using population genetic theory has shown that N < 50 in human colons and < 20 in small intestines (Chen, et al. 2019). The compartmentalization is initially proposed by Cairns to be an anti-cancer strategy such that rogue, pre-cancerous cells are confined in their own niche (Cairns 1975). Chen et al. have further shown that small N is associated with strong genetic drift which would reduce the efficacy of natural selection. As a result, the small stem cell populations cannot be easily driven to become highly proliferative but may accumulate many bad mutations in a “quasi-neutral” state (Chen et al. 2019).

With the small N in each stem cell niche, there should naturally be numerous evolutionary units. The number of stem cell niches (or crypts) in the colon is more than 10^7 in adult humans (Nguyen, et al. 2010; Baker, et al. 2014). With so many small populations evolving in parallel, it does not seem farfetched to imagine multiple occurrences of clonal expansions within an individual. Indeed, recent studies have shown that normal tissues often comprise a large number of cellular clones of various sizes (Martincorena, et al. 2015; Martincorena, et al. 2018; Lee-Six, et al. 2019; Zhu, et al. 2019). Therefore, the architecture appears to predispose tissues to multiple origins of tumors. However, DNA sequence data usually suggest a process of single origin (Parsons 2008; Thirlwell, et al. 2010; Lindberg, et al. 2013; Ling, et al. 2015; Ma, et al. 2017; Wei, et al. 2017; Cross, et al. 2018; Parsons 2018; Strandgaard, et al. 2019). This study aims to resolve this apparent contradiction.

In this study, we make a theoretical conjecture that tumorigenesis is a process of multiple origins. However, as the tumor evolves to an advanced stage, a single most proliferative clone may prevail. In this hypothesis, the *process* begins with multiple clones but the *outcome* would resemble that of a single origin. The residual independent clones may become vanishingly small but, nevertheless, contribute disproportionately to the non-neutral heterogeneity of the tumor. To test this hypothesis, the conventional practice of surveying many tumors would not be adequate. Instead, extremely deep sampling of a small number of tumors should be necessary, and perhaps even sufficient, for uncovering the residual independent clones.

### I. Theory of the progression of clonality

The number of independent cell clones would depend on the extent of tissue aberration, which could be patches of normal tissues, non-malignant growth or aggressive tumors. Even a small piece of normal tissue may harbor multiple patches of small clones (Martincorena, et al. 2015; Martincorena, et al. 2018; Lee-Six, et al. 2019; Zhu, et al. 2019). In contrast, a large tumor would often comprise a single clone of a recent origin (Gerlinger, et al. 2012; Ling, et al. 2015). This would be equivalent to the concept of selective sweep in population genetics whereby a high-fitness mutation may sweep away the lesser ones (Smith & Haigh, 1974); Hermisson & Pennings, 2005; Fay & Wu, 2001). Even multiple tumors from the same patient are often of a single origin (Cooper, 2000). The degree of cell clonality forms a continuum and the issue of single vs. multiple origins depends on the tissue (normal or cancerous) of interest.

In assessing tumor clonality, the number of evolutionary units in a given tissue is assumed to follow the Poisson distribution. Such an evolutionary unit is usually a stem-cell niche; in the colon, it would be a crypt (Snippert, et al. 2010; Kang and Shibata 2013; Baker, et al. 2014). As described above, a typical crypt in the human guts contains 20 – 50 stem cells, which are often highly clonal (Hu, et al. 2013; van der Wath, et al. 2013; Baker, et al. 2014). There are n crypts, n being tens of millions in human guts.

Let each crypt have a probability of ε to develop into a cell clone (or even tumor); thus λ = n * ε would be the expected number of independent clonal growths in a tissue. The probability that an individual has x independent clones should be f(x) = e^(-λ) λ^x^/x!.

In this model, λ = n x ε is not the same for all clonal types. The trend of clonality is presented in Fig. 1 where the X-axis shows the fitness advantage of crypts that gain advantageous mutations. Here, we adopt the mutation accumulation model of Ruan et al.(Ruan, et al. 2020) whereby crypts may occasionally accumulate many mutations in a quickened pace.

**Figure 1.**
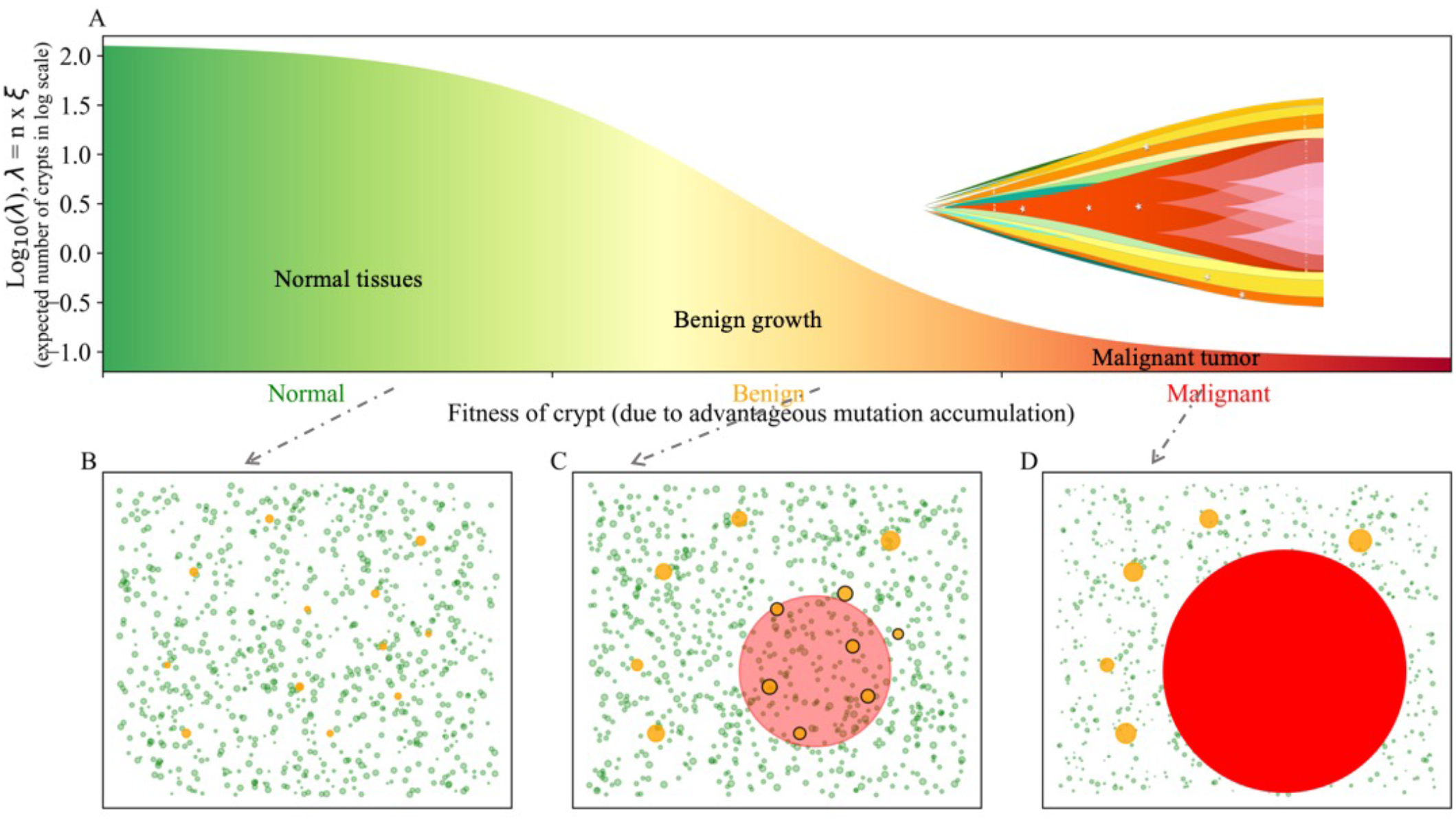
A model for the expected number of independent clones (Y-axis) as a function of the accumulation of advantageous mutations (X-axis). The X-axis is also the fitness (or the proliferative rate) of the clone. We separate the clonal expansions into 3 types: malignant tumor (red), benign polyp (orange) and non-pathological small clone (green). When sampling a patch of tissue, the possibility of detecting multiple clonal expansions is a function of mutation accumulation (Fig. B-D). Fig. 1B shows that many normal clonal expansions may occur alongside polyps (Fig. B). Fig. 1C shows that the dominant tumor clone may coexist with many smaller independent clones. As the dominant clone continues to evolve, it may drive out all other cell clones (Fig. 1D).

For the limited local expansion commonly observed in normal tissues ((Martincorena, et al. 2015; Martincorena, et al. 2018; Lee-Six, et al. 2019) λ may be in the hundreds, or even thousands (the Y axis of Fig. 1A). (Martincorena, et al. 2015; Lee-Six, et al. 2019). If we assume λ = 10 in a patch of tissue of a few cM^2 in size, then the double/single ratio would be 5 and the probability of multiple independent clones would approach 1.

For the benign growth such as polyps of the colon, λ may be slightly less than 1 for people over the age of 60 (Atkin and Saunders 2002; Oines, et al. 2017). In this group, the probability of having at least one polyp is 1-f(0) = 1 – e^(-λ) > 0.5. This means λ > 0.7 [~ - ln(0.5)] and the double/single ratio would be > λ/2 ~ 0.35. Therefore, the probability of multiple origins would be > 0.35. Fig 1B shows the most common pattern in tissues, with many small clones (green dots), as well as a few larger growths (yellow dots). Such a piece of tissue, at the incipient stage of clonal expansion, would generally be considered “normal”. Observations of Fig. 1B have been reported in many publications already (Martincorena, et al. 2015; Martincorena, et al. 2018; Lee-Six, et al. 2019).

### II. Sampling considerations for testing the theory

We now consider the advanced stages of clonal expansion toward tumorigenesis. Since the incidence for any particular type of cancer is generally < 0.1 (Feuer, et al. 1993; Merrill, et al. 1997; Svatetz 1998; Ulrich, et al. 1999; Sasieni, et al. 2011; Lloyd, et al. 2015), we assume λ < 0.1. For the colon, n is the tens of millions. Given λ = n x ε < 0.1, ε is extremely small, determined by the gradual accumulation of advantageous mutations that drive tumorigenesis. The probability of having more than 2 independent tumors should thus be negligible (Parsons 2008; Thirlwell, et al. 2010; Lindberg, et al. 2013; Ma, et al. 2017; Wei, et al. 2017; Parsons 2018; Strandgaard, et al. 2019). This is true even when multiple nodules are observed as they usually result from metastasis (Cresswell, et al. 2020; Hu, et al. 2020). Fig. 1D depicts what has been commonly reported - that the entire tumor is of a single origin.

#### Conditions necessary for observing multiple clones within the same tumor

The transition from Fig. 1B to 1D may be most interesting. Fig. 1C depicts a case of tumorigenesis when the tumor begins to expand but the area of expansion harbors several independent, albeit less proliferative, clones. Such tumors would be much more heterogeneous than tumors of single origin. It is conceivable that most tumors may comprise a dominant clone as well as many independent ones too small to detect by the standard survey. In this sense, Fig. 1C and 1D are merely different degrees of tumor progression.

In order to detect the intermediate stage of Fig. 1C with multiple clones in the same tumor, one must analyze a large number of samples (e.g., n > 100) from each individual tumor. The conventional strategy of analyzing a large number of tumors but a few samples (often one single sample) per tumor is not suited to answer the question of multi-clonal origin. Since one can only analyze a very small number of tumors by this approach, tumors have to be judiciously selected as describe below.

##### 1) Choice of tumors

Even if tumors of multiple origins are common (as long as one looks hard enough by sampling densely), a dominant clone may have sufficient time to drive out competing clones in late stage tumors (Fig. 1D). In this study, we analyze samples from a colon tumor (~ 2cM in diameter, referred to as Case I) that has metastasized to the liver. For a comparison, we choose a very large colon tumor (~ 9cM in diameter, Case II, which has also metastasized). In theory, the chance of detecting multiple clones should be very different between them. In both cases, WGS (whole-genome sequencing) is done on a subset of the samples and variants discovered in the WGS data are chosen for verification in the complete set of samples (see Methods).

##### 2) Sample number

In both tumors, very dense sampling is done as described in Fig. 2. In the main case (Case I), the primary colon tumor yields 145 micro-dissected samples (see Methods), 7 of which are subjected to whole genome sequencing. We chose 106 variants for verification among the remaining 138 samples. The same dense sampling strategy is also applied to the 4 liver metastatic lesions with 8 samples for whole genome sequencing and 147 samples for verification.

**Figure 2.**
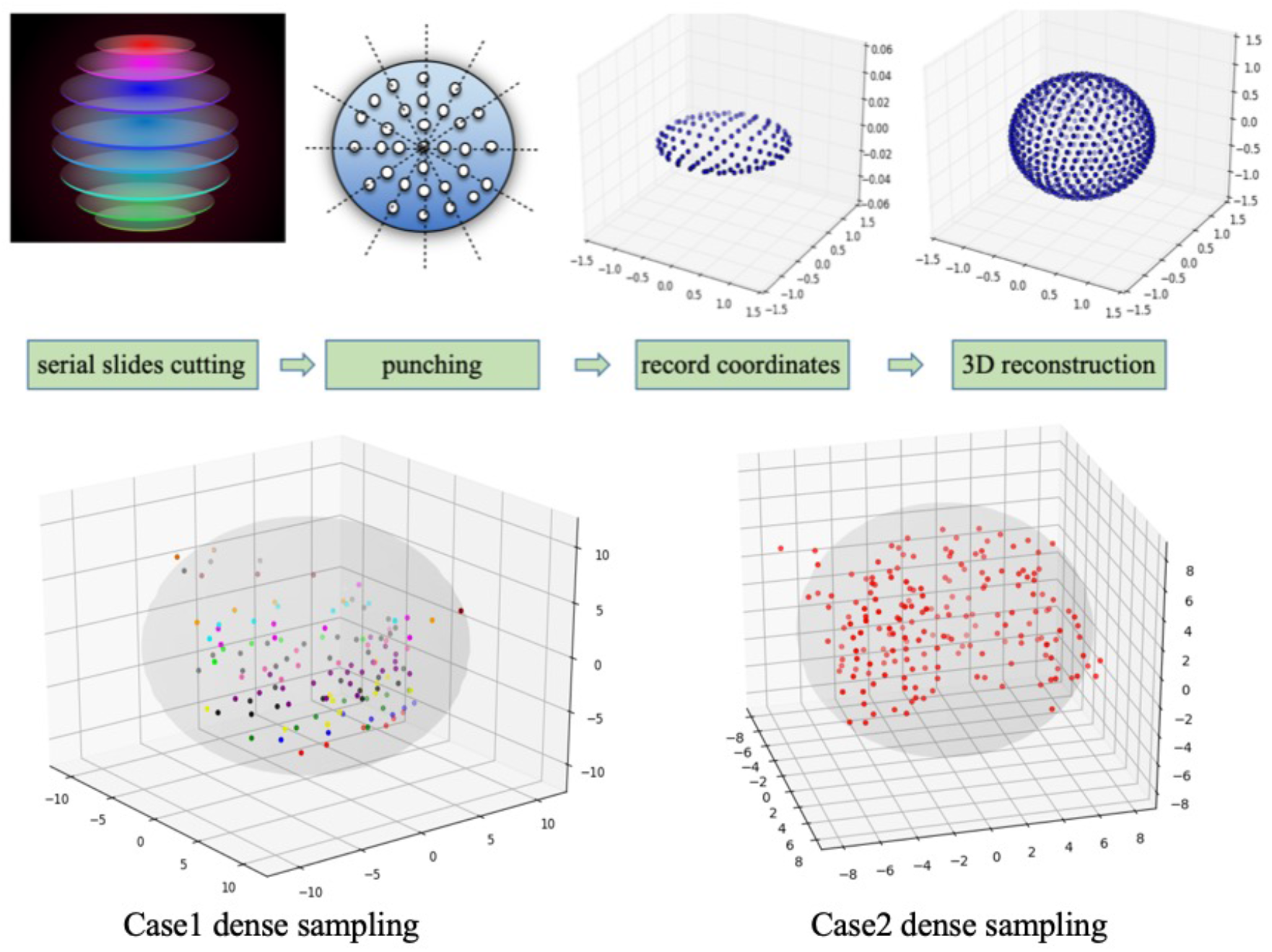
The sampling scheme in Case 1 and Case 2. (A) The tumor is sliced into 0.3mm sections using freezing microtome. Micro-dissected samples, each 0.3mm in diameter, were taken using a micropunch. Each sample cylinder contains ~3000 cells. The sampling sites are evenly distributed. The 3D coordinates of all samples were recorded for later reconstruction. (B) The spatial locations of all samples in Case1 and Case2 tumors are displayed.

In Case II, 9 WGS samples and 401 target sequencing samples (including 209 from primary tumor) were taken from the primary and metastatic tumors (colon and liver, respectively). Adjacent normal tissue as well as blood samples are sequenced by WGS as the control. (See Method and Supplements for locations of the samples)

##### 3) Sample volume

To detect minor clones, the sample volume (number of cells in a sample) has to be small. Imagine a hypothetical tumor consisting of a dominant clone accounting for 50% of the tumor as well as 50 minor ones each accounting for only 1%. In a large volume sample comprising many clones, only the dominant one could be detected while each minor clone may account for ~ 1% of the cells in the sample. In contrast, because each minor clone likely occupies a local patch of the tumor, a small-volume sample may comprise the dominant clone and, say, two minor clones that are easily detectable. This consideration may explain, at least in part, why multiple concurrent clones are not commonly observed in the TCGA data which rely on large volume samples. In this study, we do micro-dissections (see Method) such that the sample volume would be around 3000 cells for each of the many samples.

### III. Observations – Case 1 (multiple clonal origins in the same tumor)

Using the Case I colon tumor, we test the key prediction of Fig. 1 as depicted in Fig. 1C. Although most minor clones are difficult to detect, highly dense sampling (Fig. 2) may be able to reveal their presence.

#### 1) Clonal pattern of the 7 major samples revealed by WGS

The locations of the 7 samples from the primary tumor (denoted CB, C0, C1, C2, C10, C12 and C14; see Methods) subjected to WGS are shown in Fig. 3. The data, summarized in Table 1, are partitioned into groups, highlighted respectively with reddish, blue and green colors. Each row is a set of mutations that exhibit the same geographical pattern, thus forming a clone which is also named after the number of mutations defining the clone.

**Table 1.**
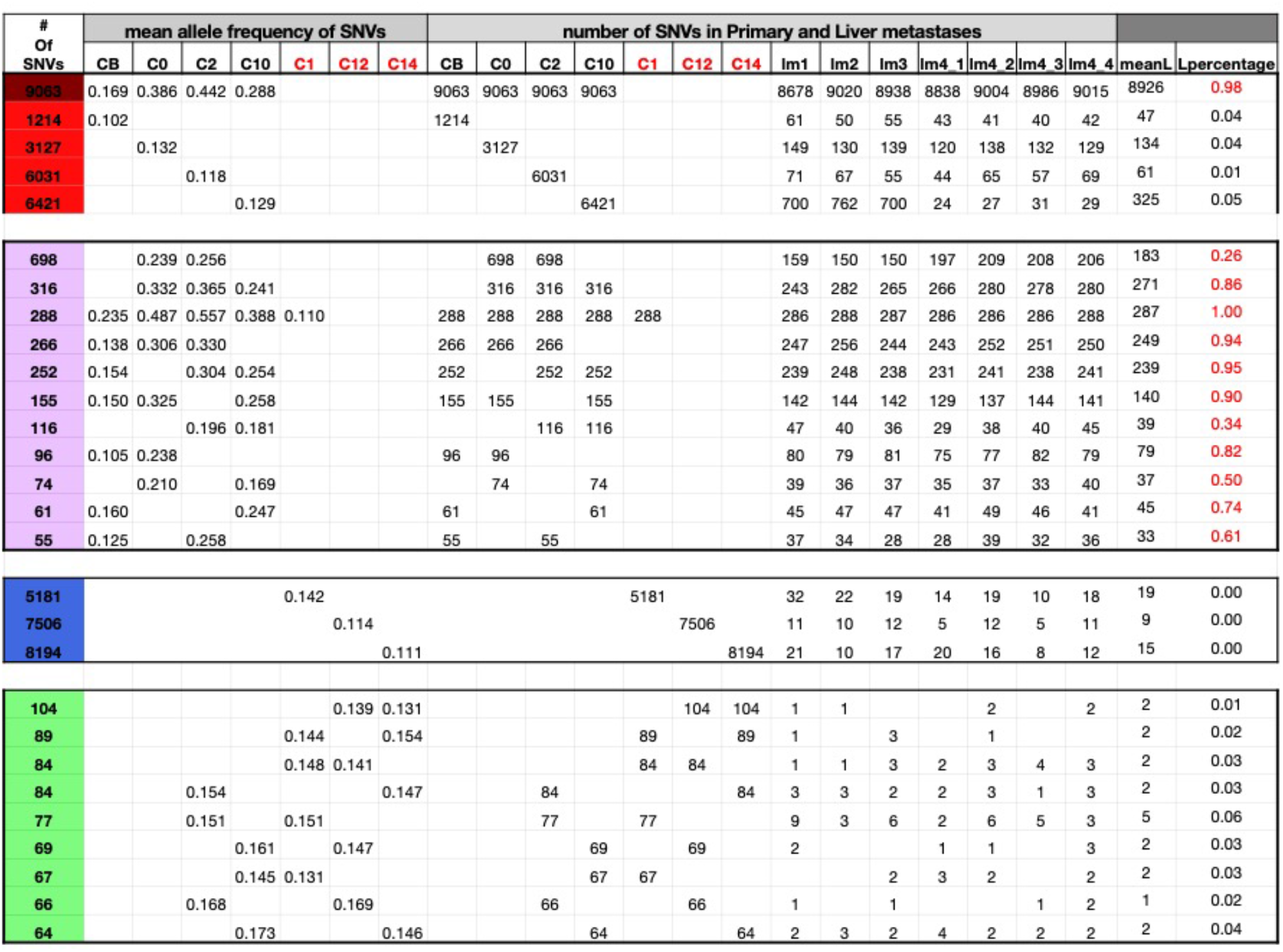
Grouping of mutations into clones in Case 1. The SNVs in the 7 WGS samples of Case1 are grouped according to the clonal patterns. Each clone, representing a group of mutations, is named by the number of mutations associated with the clone. Each row represents such a clone. Clones having similar geographical patterns are further partitioned colored categories, reddish [maroon, red and pink], blue and green. Columns of the table are divided into 3 sections: 1) Clonal frequency in the 7 WGS samples; 2) Number of mutations in these 7 samples; 3) Number of mutations in the samples from the liver metastases; and 4) [the last 2 columns] the number and proportion of the mutations observed in the liver metastases (L denoting the metastases).

**Figure 3.**
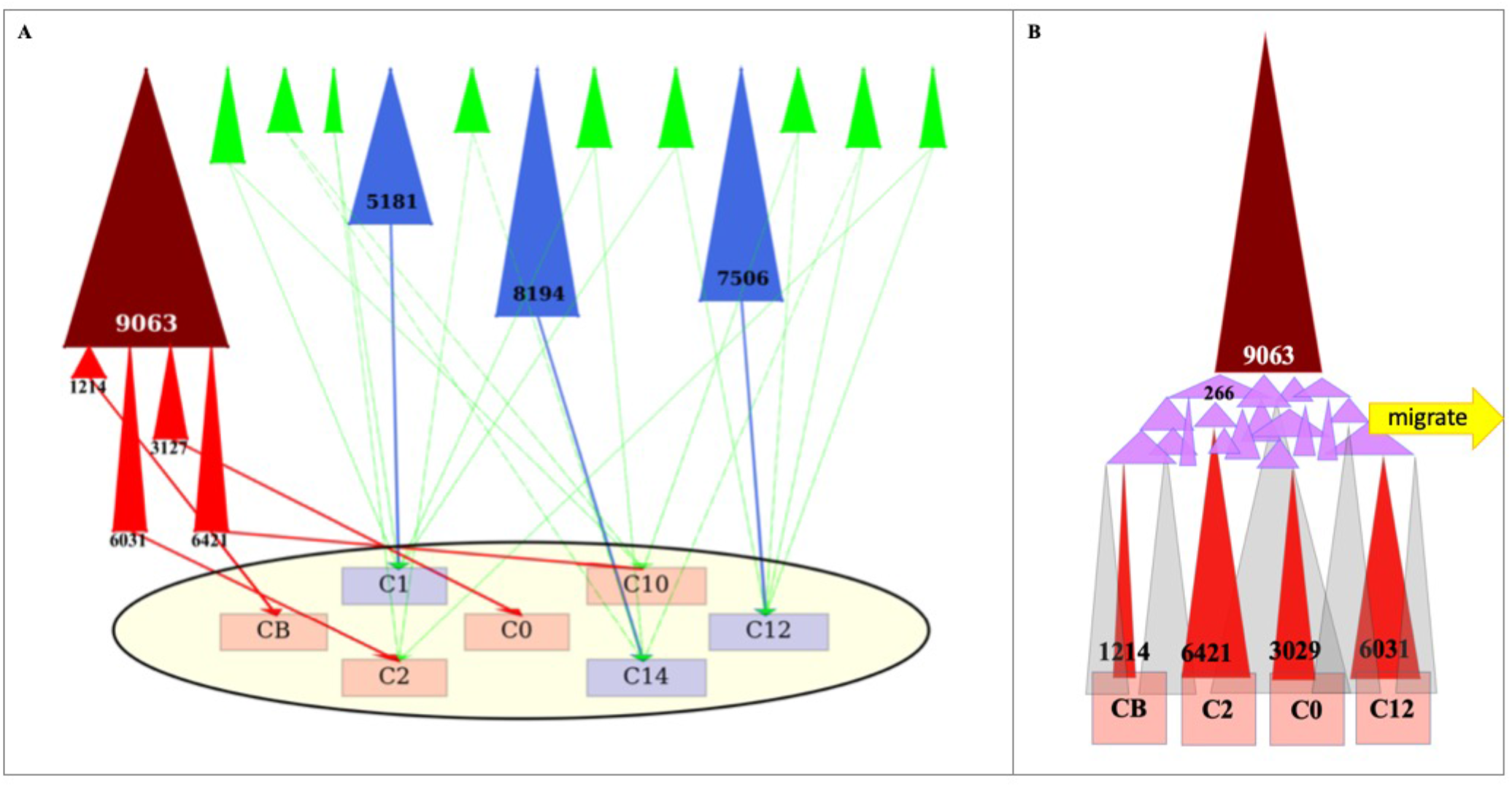
The clonal composition of the Case 1 tumor revealed by WGS (whole genome sequencing). A) Clones identified in Table 1 are represented by shaded triangles, named by the number of mutations associated with the clone. The clones are shaded in 5 different colors: maroon, red, pink (which form a family), blue and green. The distribution of these clones among the 7 WGS samples from the tumor (the oval) are shown by the thin lines. The 4 red clones are all subclones of the 9063 parent clone. Note the presence of 13 independent clones consisting of a dominant clone (represented by the maroon and its red sub-clones), 3 large blue clones and 9 small green clones (see Table 1). Cells of the small green clones are dispersed, presumably by the dominant maroon clone as it expands. B) The widely distributed maroon clone spawns the 4 sample-specific red clones via a large number of pink clones, which represent a stage of geographical expansion of the maroon clone. Hence, the pink clones are less localized than the larger red subclones. The approximate timing of the metastasis to the liver is indicated by the block yellow arrow at the pink clone stage.

The reddish group in Table 1 has two sub-groups designated by red (Group 1) and pink (Group 2) colors. These 2 groups of clones are all found in the core samples of CB, C0, C2, C10 (also colored reddish in Fig. 3). Group 3 (colored blue) clones are found in C1, C12 and C14 samples while Group 4 (colored green) are scattered among the seven samples. The distinction of these four groups is informative about the evolutionary dynamics of the tumors.

For Group 1, the 9063 clone is the most dominant clone in size and depth (i.e., number of mutations). Group 1 comprises, in addition to the 9063 clone, 4 other clones that are specific to each of the 4 core samples, CB, C0, C2, C10. These clones (Clone 1214, 3127, 6031 and 6421) are all subclones of Clone 9063 as depicted in Fig. 3 and Table 1. A subclone has all the mutations of the parent clone (in this case, the 9063 mutations) as well as additional ones. These 5 red clones listed on the top of Table 1 are the main clones that dominate the primary tumor. Group 2 clones (colored pink) are also found in the four samples (CB, C0, C2, C10. However, they have far fewer mutations than Group 1 clones and are found in more than one sample. These clones will shed light on the evolution of Group 1 clones, as described in the next section.

Interestingly, 3 of the 7 samples (C1, C12, C14) do not harbor any mutations of the Group 1 or Group 2 clones. Each of the three samples, instead, is dominated by an independent minor clone of Group 3 (Clones 5181, 7506 and 8194; colored in blue). Therefore, the primary tumor depicted in Fig. 3 comprises at least 4 large clones. They are the 9063 clone (plus its 4 subclones in Group 1; all colored red) and the 3 blue-colored clones (Group 3). Furthermore, the clones are scattered widely within the tumor. For example, the clone in C1 is surrounded on all sides by the main clones and are distantly located from other Group 2 clones. The depiction in Fig. 3 thus supports the conjecture of Fig. 1C showing smaller independent clones engulfed by a dominant one.

The last group in Table 1 is the Group 4 clones, colored green. As depicted in Fig. 3, they usually straddle two samples. Group 4 clones are likely remnants of the clonal expansions commonly observed in normal tissues ((Martincorena, et al. 2015; Martincorena, et al. 2018; Lee-Six, et al. 2019)). As can be seen in Table 1, there are 9 such normal expansions (see also Fig. 3). It is worth noting that these clonal expansions are detected when the cells are found in different samples. For example, Clone 84 are found in C12 and C1 and Clone 66 are present in C12 and C2. Hence, there are likely numerous clonal expansions that escape detection because the cells are not dispersed. In short, the primary tumor is composed of at least 13 clones - one large, 3 medium-sized and at least 9 small clones.

The dispersal of the small “normal” clones is instructive about cellular movement within the same tumors. As the tumor grows and various cell clones expand in size, cells may be passively pushed around, resulting in the scattering of the same clone in multiple locations. Such passive dispersal should not be considered true cell motility, an issue to be addressed later by the analysis of large clones. Cell motility could be an important factor in the outcome of clonal competition (and, hence, the presence/absence of multiple independent clones).

#### 2) Clonal pattern of the 138 validation samples (revealed by 106 SNVs)

In Fig. 3, the clonal origins are depicted in only 7 samples from a slice of the tumor. Here, we genotype 138 samples from the primary tumor as shown in Fig. 2 In total, 106 SNVs, identified by WGS mainly in the Group 1 and 2 clones, are used to genotype the 138 samples. The clonal patterns are shown in Fig. 4 where the sample locations are displayed in the top row of the heat map. The black and grey bars denote, respectively, samples from the left and right side of the tumor. The orange bar denotes the metastatic samples while the cyan bar denotes either blood or non-cancerous tissues adjacent to the tumor. Corresponding to the heatmap of Fig. 4 is the 3D distribution of the clones displayed in Fig. 5. In this figure, the tumor is portrayed in 9 panels, each of which is a composite of 2-3 neighboring slices from the 18 slices of the tumor. Roughly between the 4^th^ and 5^th^ panel is the pattern of Fig. 5. The 3D distribution of each of the 5 clones in Group 1 is portrayed by a group of 9 panels, described below.

**Figure 4.**
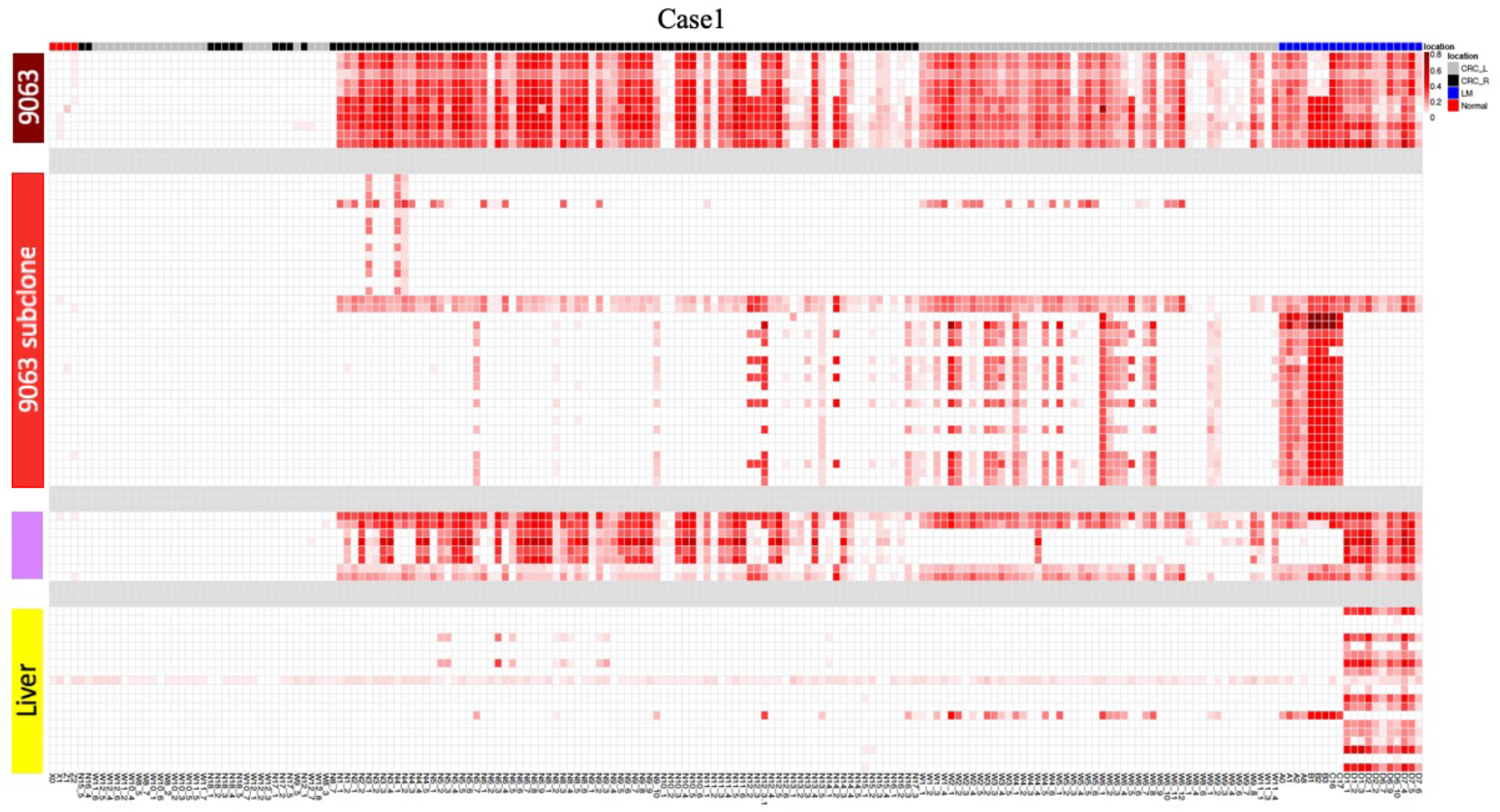
Mapping the distribution of the maroon, red and pink clones across the 138 samples from the primary tumor and the liver metastases. Each column represents a sample. labeled by different colors (black, orange and cyan) on the top row of the heatmap to show their locations. Each row represents the distribution of an SNV across the 138 samples. The collection of 106 SNVs are separated into 4 groups (Maroon, red and pink as shown in Fig.3 plus the yellow-color group, representing mutations of the liver metastasis). The color intensity in each box indicates the frequency of the SNV in each sample.

**Figure 5.**
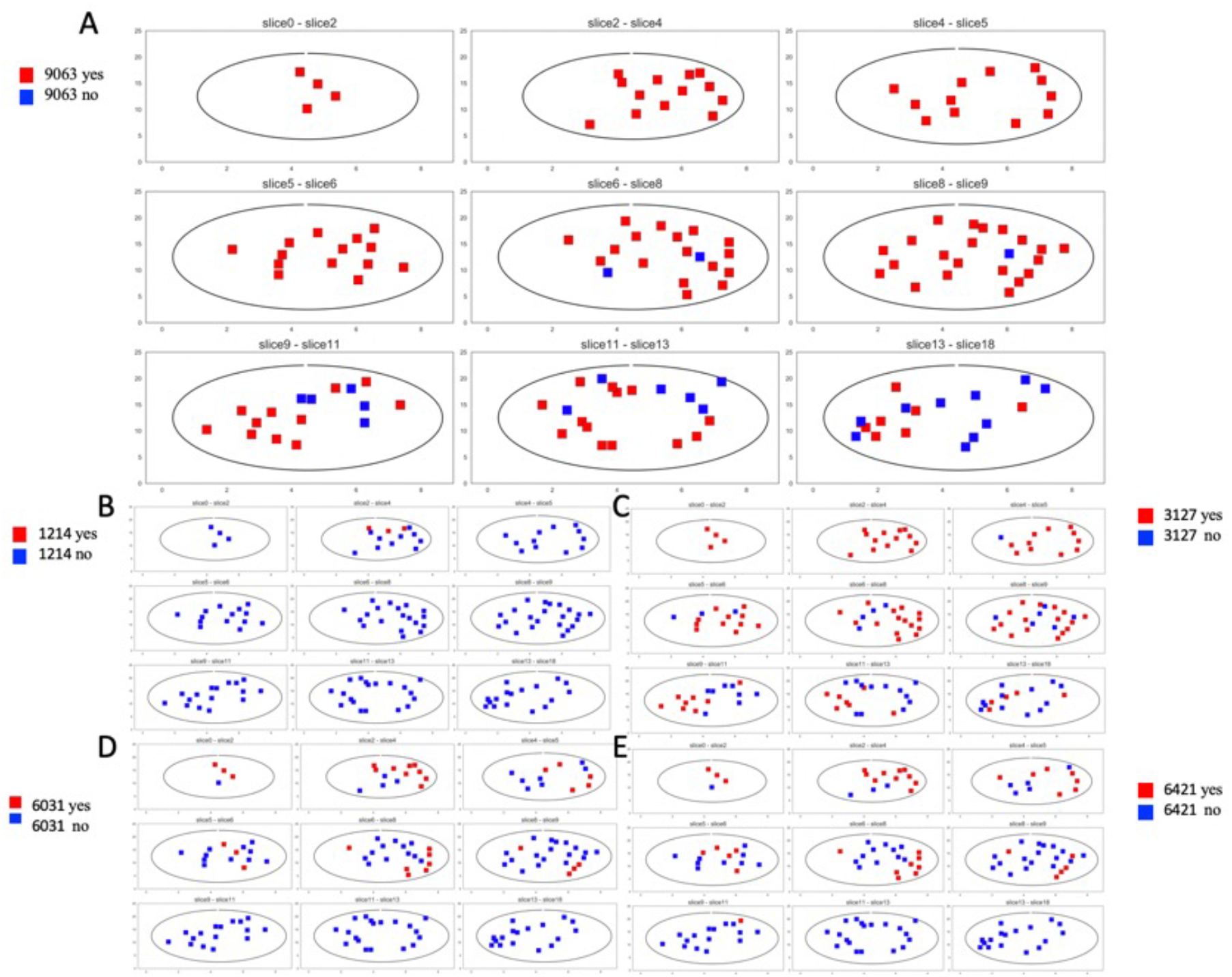
The 3D distributions of SNVs across 138 samples from 9 serial planes in Case 1. Each of the five groups of mutations is shown as present (red) or absent (blue) among the 9 planes (one plane for each panel) of the tumor. Each plane represents a composite of 2-3 neighboring slices from the 18 serial slices of the tumor as illustrated in Fig.2. Note that the SNVs from the major 9063 clone (maroon color in Fig. 3) are not found in many samples of the lower 3 planes. The geographical distribution of SNVs from the 4 red clones (the subclones of the 9063 parental clone) are showed from Fig, 5B to Fig. 5E. These subclones are indeed strongly localized, supporting the interpretation given in Fig. 3.

**Figure 6.**
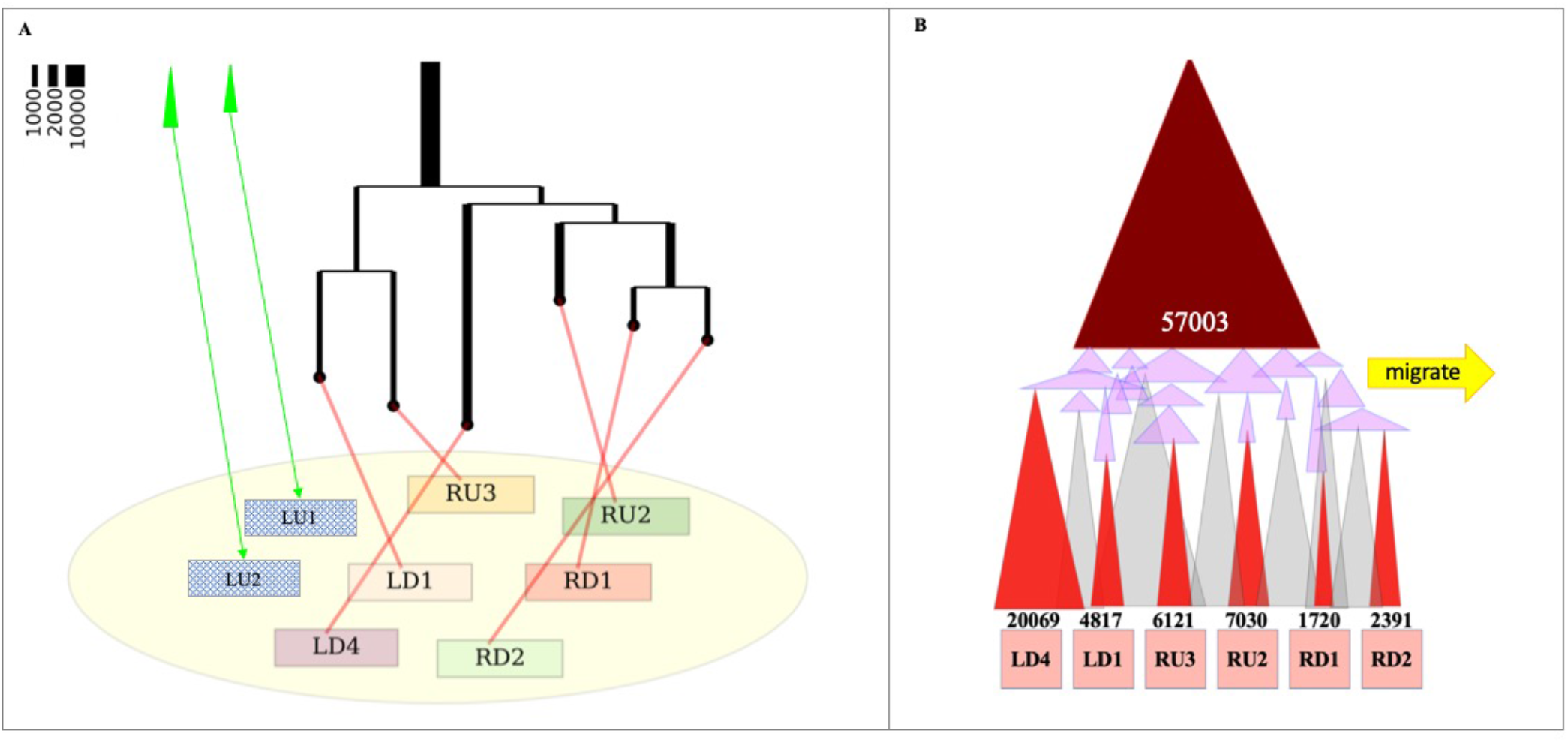
The clonal composition of the Case 2 tumor revealed by WGS. See the descriptions of Fig. 3, which apply to Fig. 6 as well.

The 9063 clone – In Table 1, one can see clearly that the 9063 clones indeed populates 80% of the samples all over the tumor as well as the entire metastases. Not all the entire set of 9063 mutations are distributed equally to all parts of the primary or the metastatic tumors. Some mutations that occurred late did not ride with the earlier waves of spread, as seen in the much lighter squares in the heatmap. The key message is that, while this clone has successfully metastasized to the liver, it still fails to displace some of the minor clones in the primary tumor.

The 3D distribution of the 9063 clone is shown in Fig. 5A. This clone occupies the top portion of the tumor completely but has not reached the lower portion of the primary tumor. Fig. 3(3A?), being a slice in the middle, happens to present the average prevalence of the 9063 clone. This main clone shows contiguity in its spread, as can be reconstructed from the 18 slices. In short, cells move *passively* as the tumor mass expands. This will be a recurring theme.

The 4 subclones in Group 1 – Note that the subclones (red-color clones of Table 1) are all specific to each of the 4 samples shown in Fig. 3. Within the entire tumor (see Fig. 5B-4E, each having 9 panels), they exhibit a range of characteristics. The 3127 subclone of Fig. 5C is the most broadly distributed but it still occurs in less than 50% of the samples (note that the presence of the clone is shown with a red dot even at a very low frequency). The remaining 3 subclones are all rather locally distributed, this being particularly evident for the 1214 clone.

We interpret the mutations of these subclones to have emerged rather late during the tumor growth. At the late stages, cells do not get pushed out on the expanding edge and thus result in the much more highly localized distributions. The number of mutations each clone accrue should also be a measure of the passage of time. In short, most of the cells of these clones are constrained in a localized area for a substantially long period of time. This is another indication that cells of the expanding clones do not actually migrate; they move only as a function of the tumor growth.

#### 3) The detailed evolutionary pattern of Group 1 clones (revealed by Group 2; see Fig. 3B)

Clones defined by Group 2 mutations (colored pink in Table 1) have the depth of 50 - 700 mutations. They are found in 2 – 3 major samples (CB, C0, C2, C10). With the sole exception of the 288 clone, they are distributed in a subset of samples of the 9063 clone (see Table 1). A detailed picture is seen more clearly in Fig. 4 across all 138 samples (plus those in the metastases).

To understand Group 2 mutations in relation to the evolution of the primary tumor, we provide a more realistic portrait of the clonal expansion (see Fig. 3B; vis-a-vis Fig. 3A). In Fig. 3B, a series subclonal expansions are shown to have occurred between the 9063 clone and its sample-specific sub-clones (i.e., the 6421 group). Group 2 mutations defined this group of intermediate clones (pink color). Therefore, the oldest clone (i.e., 9063) has the widest distribution and the 6421 group of subclones are the youngest as well as the most spatially constrained. Group 2 clones have a spatial distribution in between them since their ages are between the two sets of major clones. For example, the 266 clone as marked can be found in the CB, C0 and C2 samples. In the same framework, each sample comprises more than one clone. Those clones colored in light gray are hypothetical clones, the mutations of which are not identified.

Fig. 3B depicts clonal movement solely as a consequence of clonal expansion and inter-clonal crowding. The overall pattern may thus resemble the “glacier movement”. In this mode, older clones spread more widely and the youngest clones are highly localized as conceptualized in Fig. 3 and observed in Fig. 5. Whether the low cell motility contributes to the multi-origins of the primary tumor will be addressed in Discussion.

#### 4) The continual dominance of one single clone (revealed in the liver metastases)

Extending the observations presented in Fig. 3A and 2B using the model of Fig. 1, we may predict the “future” of the primary tumor. As stated, four metastases in the liver have been observed, sampled and sequenced. The clonal frequencies are given in Table 1 under LMi (for liver metastasis 1 – 4). The last column shows the proportion of the mutations of each clone in the LMi samples. This column is most informative about the growing dominance of some clones in the next phase of the evolution of the primary tumor.

It is clear that the metastases originate from the 9063 clone, since 98% of its mutations are found in the liver samples. The second prevalent group is Group 2 clones. The proportion of the mutations from this group present in the metastases ranges between 26% to 100% with an average of 72%. Interestingly, the 4 subclones of 9063 in Group 1 (the red-color ones) are rarely present in the metastases with only 1% - 5% of the mutations there. This pattern permits us to pinpoint the timing of metastasis to be during the expansion of the pink clones in Fig. 3B, before the red clones begin to proliferate. One does not expect the growth of Group 3 or Group 4 clones which appear localized with limited proliferative capacity. Indeed, Table 2 shows their near absence in the metastases.

**Table 2.**
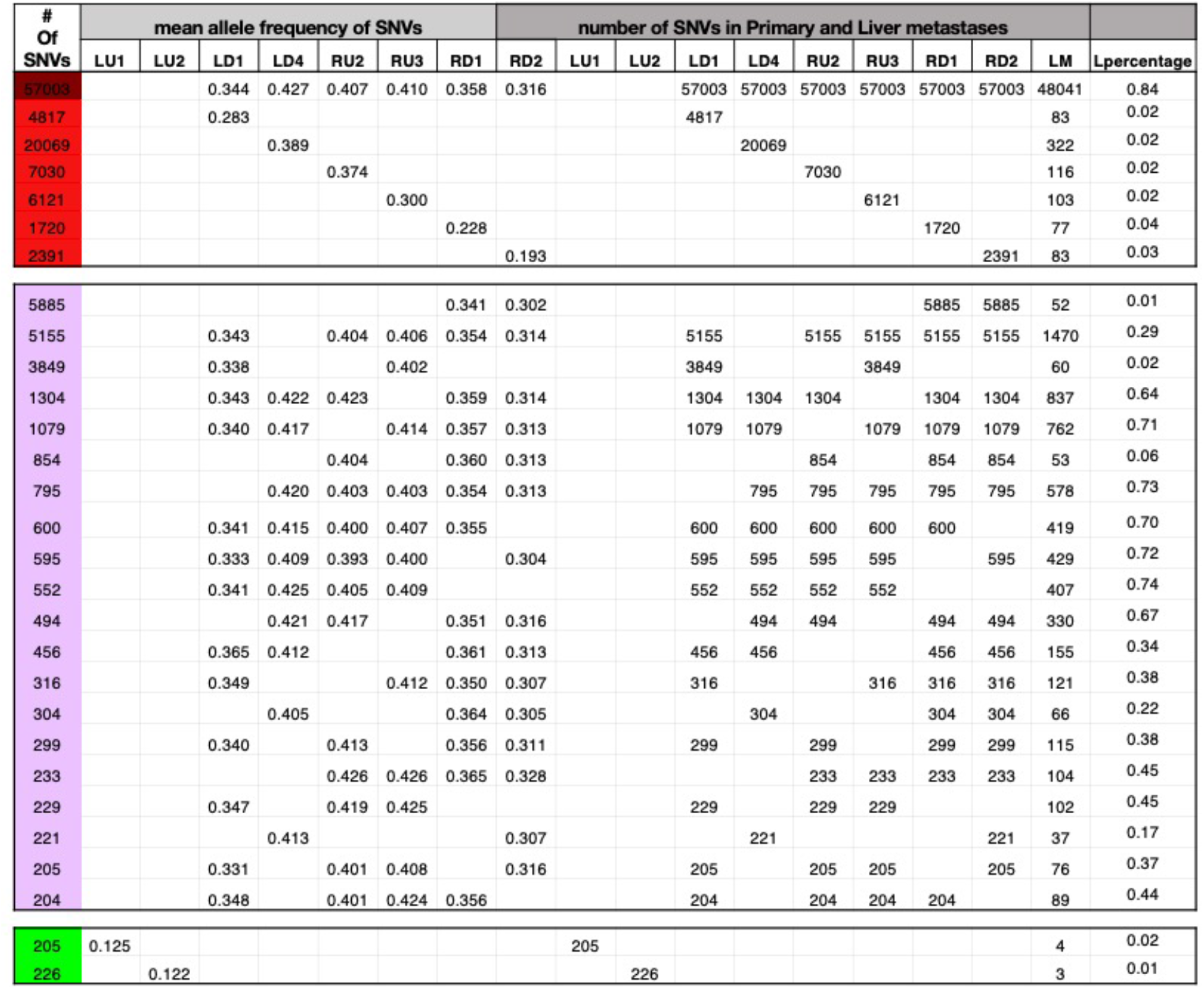
Grouping of mutations into clones in Case 2. See Table 1 section for details.

### IV. Observations – Case 2 (The advanced stage with one dominant clone remaining)

In the scheme of Fig. 1, tumors of multiple clonal origins should not be uncommon, although the sampling may have to be nearly exhaustive to reveal the multi-clonality. By extension, at the very advanced stage, clonal competition may reduce the field to a single dominant clone. Clonal competition may be analogous to the evolution of warring states in the human history as the emerging empire engulfed smaller city-states along the way.

In the second case of a large colon tumor (~ 7 cM in the longest dimension, which has one liver metastatic lesion), we ask i) whether the primary tumor has a simple clonal composition with a single dominant clone; ii) whether remnants of minor clones could still be found; and iii) how the presence/absence of the minor clones may inform about the process of clonal competition. Here, we are searching for very rare minor clones in order to test the hypothesis of Fig. 1. In this case, 209 samples are dissected from the primary tumor including 9 samples used for WGS. Then, 64 variants are validated in all the other samples.

Among 8 WGS samples of Case2, all but two samples (LU1 and LU2) are derived from a dominant clone which harbors 57003 mutations (see Figure 5 and Table 2). However, the 57003 clone is far more prevalent than the WGS survey might indicate. The target validation for 209 samples demonstrates the similar high prevalence of the major clone as the clonal mutations detected in 98% of the validation samples (Figure 7). This scenario, consistent with Fig 1D where a single clone reigning the whole tumor, is the most widely depicted pattern of cancer evolution(Ling, et al. 2015). Besides the 57003 old clonal mutations (dark red), the other group I and II mutations (red and pink clones) representing later local expansion can also be found in Case2 (see Fig. 5B).

**Figure 7.**
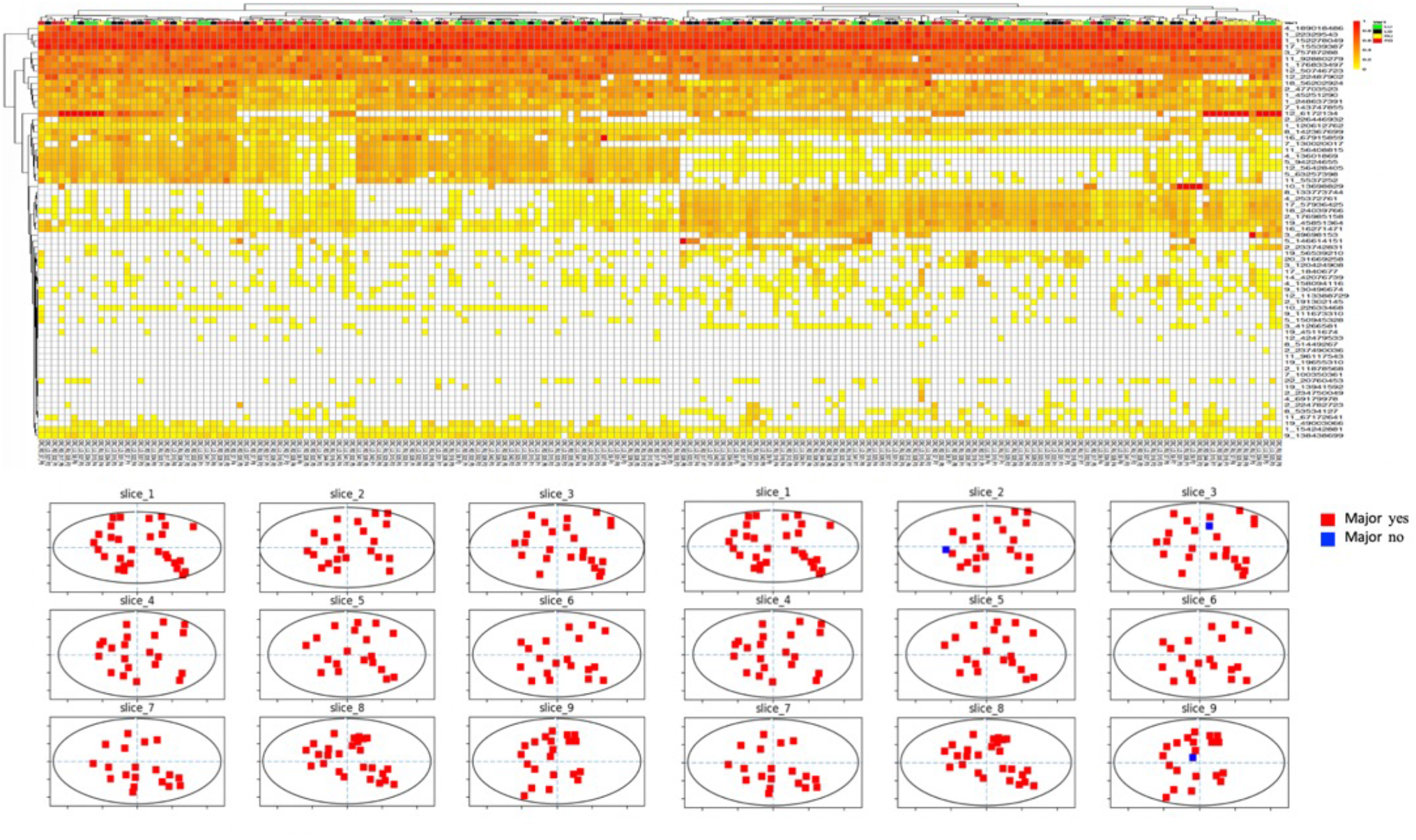
The geographical distributions of SNVs across the samples of Case 2. See the descriptions of Figs. 4-5, which apply to Fig. 7 as well.

In comparison with Case 1, Case 2 is distinguished by 1) the dominant 57003 clone is present in 98% of the samples with LU1 and LU2 being the rare exceptions; 2) the absence of many small independent blue-colored clones; 3) few green-colored clones can be detected. These patterns suggest the shrinking of the minor clones when the dominant clone expanded. All of above observations indicate the dominant clone already prevail over the other independent clones and take over the entire space when the tumor evolve to the late stage.

In the liver metastatic tumor of Case 2, 84% of the 57003 clonal mutation as well as 6%-74% of the pink-colored mutations are present. While for the group I red color mutations (4817,20069, 7030, 6121,1720 and 2391), only ~ 2% of mutations can be found in Liver metastatic samples. It is clear that the timing of metastasis is similar to Case1. Cell migration appears to happen when the dominant clone begins to expand locally (indicated by the pink color). In short, Cases 1 and 2 are qualitatively similar. Case 1 shows all the signs of a tumor in the process of crowding out the competitors whereas Case 2, a much larger primary tumor, has accomplished much of the actions. Note that their metastases show similar clonal compositions.

## Discussion

The existence of genetic diversity within the same tumor has been extensively reported and discussed(Ling, et al. 2015; Tao, et al. 2015; Wang, et al. 2018; Wen, et al. 2018). Almost all the genetic diversities are generated during the process of the clonal expansion from a single progenitor cell. Hence, much of the genetic diversity is shared. The debate often emerges on the fitness differences imparted by the mutations. The debate on cancer evolution therefore parallels the neutralism-selectionism debate on organismal evolution (Wu et al, 2016).

In a single clonal expansion reported in the literature, it is plausible that the many closely-related subclones are neutral in fitness and functionally equivalent (Ling, et al. 2015; Sottoriva, et al. 2015). In this study, we focus on a different kind of genetic diversity. The independent clonal expansions accrue entirely different sets of mutations. Thus, when two clones emerge independently, each carrying its own >10,000 mutations, these clones would seem unlikely to be functionally equivalent.

Can functionally divergent clones coexist in the same tumor? In cancer genomics, the fitness of a clone is usually equated to its ability to proliferate (many many references). In this view, the performance of the tumor would depend on the most highly proliferative clone in a “winner takes all” scenario. This overly simplistic view of fitness does not permit functional heterogeneity. However, in nature, we do not consider grasses to be more fit than woody plants, or rodents to be more fit than tigers. Likewise, in somatic evolution, the coexistence of divergent clones with different proliferative abilities can often be found in colon tumors with the cancer-adjacent polyps (CAPs, cite Lisa Boardman’s papers on CAPs). These CAPs often harbor more mutations than the neighboring tumors, suggesting that they are not merely less-evolved.

Li et al. (Li, et al. 2020) show that cells of tumors may follow the life-history strategies in organismal evolution. Given the limited amount of resources for allocation to reproduction vs. competition, a species may choose to produce many progenies with low competitiveness (like grasses) or fewer progeny that are highly competitive (like woody plants). The choices are referred to, respectively, as the r-strategy (grass) or K-strategy (woody plant) based on the logistic equation of population growth. In the equation of dN/dt = rN [K-N]/K, r is the Malthusian parameter of population growth and K is the carrying capacity of the population in the given environment. Li et al (Li, et al. 2020) show that cancer cells may evolve toward the r-strategy on the periphery of the tumor whereas the K-strategy is favored in the center of the tumor.

In this light, the conventional view of “the more, the better” in tumorigenesis may be too restrictive. Alternatively, the “minor” clones of Fig. 3 might be K-strategists, slowly proliferating but highly competitive. The proliferative clone may only be able to go around the K-patches, rather than to eliminate them. The two types of cells occupy different ecological niches. In a companion study, we suggest that CAPs may be the K-type clones, rather than the precancerous tissues. A clinical implication is that target gene-therapy may eliminate the r-clones leaving K-clones to grow slowly. K-clones are hence the reservoir of drug-resistance cells. Subsequently, some of the K-cells may acquire the characteristics of the r-cells to become rapidly proliferating. There is a time delay in the switching of strategies, which may explain tumor recurrences after a time lapse.

In this analysis, it is assumed that tissues are homogeneous in providing a micro-environment for clonal expansion. However, the tissues may be patchy with some micro-environments being particularly suited to clonal expansion. Factors may include mutation rate elevation, inaccessibility to immune cells, blood supplies and a host of other factors (many references). In that case, multiple clones observed in Fig. 3 would be common since the presence of a tumor is indicative of a fertile micro-environment for clonal expansion. In conclusion, independent origins of clonal expansion greatly enhance the functional diversity within tumors. We show that very deep and broad sampling may often uncover such patterns. This sort of diversity could be functionally more significant than the diversities generated from a single clonal expansion.

## Methods and supplementary

### Patients information

The Case1 patient was a 70-year old female, with a preoperative diagnosis of sigmoid colon moderately differentiated adenocarcinoma with liver metastasis. The patient refused to receive any other treatment except for surgery. She eventually underwent laparoscopic sigmoidectomy. During the operation, 4 liver lesions were identified and resected, all of which were confirmed to be metastatic liver adenocarcinoma. The postoperative pathology showed that her sigmoid cancer was a 22*20mm protuberant mass, and her pathological TNM stage was T2N1cM1. The Case2 patient was a 35-year old female. Her preoperative diagnosis was poorly differentiated adenocarcinoma located at the descending colon with liver metastasis. The liver lesion was 21*19mm in size and on the left lobe of liver. She underwent a radical descending colectomy, and the liver lesion was also resected. Her descending colon cancer was shown to be a 70*55mm ulcerative mass. The postoperative pathology confirmed the diagnosis of descending colon adenocarcinoma with liver metastasis, with a TNM stage of T3N0M1. Informed consent was obtained from both of the two patients.

### Dense sampling strategy of primary and metastatic tumors

We took paired normal and primary colorectal as well as liver metastatic tumors from the two patients and sliced them into a series of contiguous frozen sections of 0.3mm thick with the Leica freezing microtome after embedding. Micro-dissected samples of 0.3mm in diameter were taken with a micro-punch of 0.3mm inner diameter. Each cylinder sample contains ~3000 cells (Fig S1). The coordinates of all samples were recorded for later 3D reconstruction.

In Case1, 15 (for WGS)+ 253(for Target sequencing) micro-dissected samples were taken from the tumors, among which 7(WGS)+138(TS) are from primary tumor while others are form liver metastatic lesions. 9 WGS samples and 401 target sequencing samples (including 209 from primary tumor) were taken from tumors of case2. Adjacent normal tissue as well as blood samples were also taken for control. The x,y,z coordinates of all samples are recorded in Table S1. Genomic DNA was extracted using Tiangen Micro DNA kit and then subjected to sonication using Covaris. The library was built with VAHTS TM Universal DNA Library Prep Kit for Illumina^®^ (ND604, Novizan).

### Sequencing data processing

After whole genome sequencing, we filtered out low-quality reads and mapped raw sequencing reads to GRCh37 human genome with BWA. The filtered reads were then processed through GATK and we then used mutect2 and varscan to detect SNVs and filtered out the reads of: 1) prevalent human SNPs; 2) sequencing depth <10X; 3) mutated reads number <5 or 4) more than 1 SNVs coexisting in 1000bp length.

### Polymorphic mutations selection and target sequencing

Variants of different groups of mutations (Table 1) discovered in the subset of WGS samples are chosen for verification in the complete set of micro-dissected samples. In case1, 106 SNVs are used to genotype the 138 primary samples as well as 115 liver metastatic samples. As the tumor size of case2 is larger than the tumor in case1, more samples (209) dissected from the primary and 200 from liver metastasis are used for validation of 64 variants identified by WGS data.

